# The role of age in the relationship between brain structure and cognition: moderator or confound?

**DOI:** 10.1101/2025.05.19.654872

**Authors:** Ben Griffin, Chetan Gohil, Mark Woolrich, Stephen Smith, Diego Vidaurre

## Abstract

Understanding how differences in brain structure relate to differences in cognition across the lifespan is essential for addressing age-related cognitive decline. Since age is strongly associated with both brain structure and cognition, predictive models often risk simply capturing age effects. To mitigate this risk, deconfounding is typically applied to remove the effects of age. Here, we propose to treat age instead as a moderator variable, therefore capturing changes in how brain structure and cognitive abilities are statistically connected. For this view to hold, variations in brain structure linked to differences in cognitive performance in older subjects (e.g., related to disease) would differ from those in younger subjects. Using structural brain imaging data from the UK Biobank we found an asymmetry in generalisability: models trained on younger subjects successfully predicted cognition in older subjects, but models trained on older subjects failed to generalise to younger individuals. These findings reveal a trade-off between model specificity and generalisability, suggesting the optimal approach—whether age-specific or pooled—depends on the research or clinical goal for the target population.

## 1 Introduction

Identifying robust associations between brain structure and cognition is critical for both basic neuroscience and clinical applications. Predictive models that reveal relationships between brain data (e.g., structural MRI) and cognitive abilities (e.g., fluid intelligence) are important for uncovering these associations, improving our understanding of cognitive decline and neurological disorders such as Alzheimer’s disease (Solé-Padullés et al., 2009) and other dementias (Suzuki et al., 2019).

Capturing these brain-cognition relationships is complicated by the fact that ageing affects both brain structure and cognition in complex ways that vary across individuals. For instance, different cognitive abilities show different relationships with age (Horn & Cattell, 1967); crystallised abilities generally improve until 60 before plateauing, whereas fluid abilities begin to decline from age 20 (Murman, 2015; Salthouse, 2010). Similarly, different brain regions show distinct patterns of age-related atrophy (Bethlehem et al., 2022; Fjell & Walhovd, 2010). The complexity increases when considering structure and cognition together. For example, dividing adults aged 18-88 into younger and older groups revealed a weaker relationship between white matter and certain cognitive functions in older adults, suggesting that some brain-cognition associations become less stable with age (de Mooij et al., 2018).

These complexities make robust predictive modelling challenging, particularly when accounting for age. Although age is a key variable in certain studies of age-related conditions (Davis et al., 2018; Jack et al., 2019), it is often treated as a confound, and typically linearly regressed out from both the brain data and cognitive variables (Azevedo et al., 2023; Bracher-Smith et al., 2022). This approach ensures that identified relationships are truly reflective of brain-cognition associations independent of age, at least under assumptions of linearity. Nonetheless, models can still show age-related bias. While such biases are well documented for sex, ethnicity, and socio-economic status (Li et al., 2022; Martin et al., 2019; Obermeyer et al., 2019), they also arise for age (Greene et al., 2022; Wu et al., 2023). This is consistent with evidence that predictive models can also show limited generalisability across behavioural domains (e.g., cognition vs. personality; Chen et al., 2022). These complexities highlight the importance of developing predictive approaches that perform robustly across age groups and call into question whether age should be treated merely as a nuisance variable to be removed.

Therefore, we move beyond the treatment of age as a linear confound. We address whether the nature of the brain-cognition relationship is consistent across age groups, or if, on the contrary, age *moderates* this association—meaning the strength or even the direction of the relationship itself changes depending on age. While some studies test moderation using interaction terms, these approaches may fail to distinguish between true interaction effects and underlying nonlinear relationships (Rimpler et al., 2024). Here, instead, we test generalisation using age-stratified predictive models. Given that ageing involves different underlying mechanisms (e.g., neurodegeneration), we hypothesise that the structural brain features associated with cognitive performance in older adults may differ from those in younger populations.

Using data from UK Biobank (UKB) (Sudlow et al., 2015), we found a difference in how predictive models generalise across age. While certain brain-cognition associations appear stable across the lifespan, we reveal an asymmetry: models trained on younger subjects generalise well to older subjects, but models trained on older subjects fail to capture the relationships relevant to a younger population.This suggests that while certain brain-cognition associations are stable across the lifespan, there may be increased noise, variability, or complexity in the relationships for older subjects that impacts model generalisability. These results highlight a trade-off between specificity and generalisability, with important implications for both research and clinical applications.

## 2 Methods

### 2.1 UKB neuroimaging and non-imaging data

We analysed data from UKB, a large-scale neuroimaging dataset that includes subjects aged 47 to 83 years. Our study explored the relationship between a composite cognitive measure, specifically the first principal component of the top 30 cognitive traits selected from the 1,331 available traits in UKB (see details in **Subsection 2.1.2**), and 1,439 structural IDPs. These IDPs comprised 1,436 T1-weighted MRI measures, including Regional and Tissue Volume, Cortical Area, Cortical Thickness, Regional and Tissue Intensity, Cortical Grey-White Contrast, and White Matter Hyperintensity Volumes, along with three T2-weighted MRI measures. Our analysis included 25,170 UKB subjects, after excluding those with missing values for any of the 30 cognitive traits or for age. Missing values in structural IDPs were imputed using the mean to maintain a larger sample size. **Figure 1a** shows the age distribution of the UKB cohort, split into younger and older halves at the median age to create equally sized groups. This division was used in our multivariate prediction analyses to balance sample size and stability. **Supplementary Figure SI-1** further splits these halves by sex, confirming similar male-female distributions within each age group. For our univariate analyses, subjects were instead stratified into quartiles to capture finer age-dependent variation (**Supplementary Figure SI-2**).

**Figure 1.**
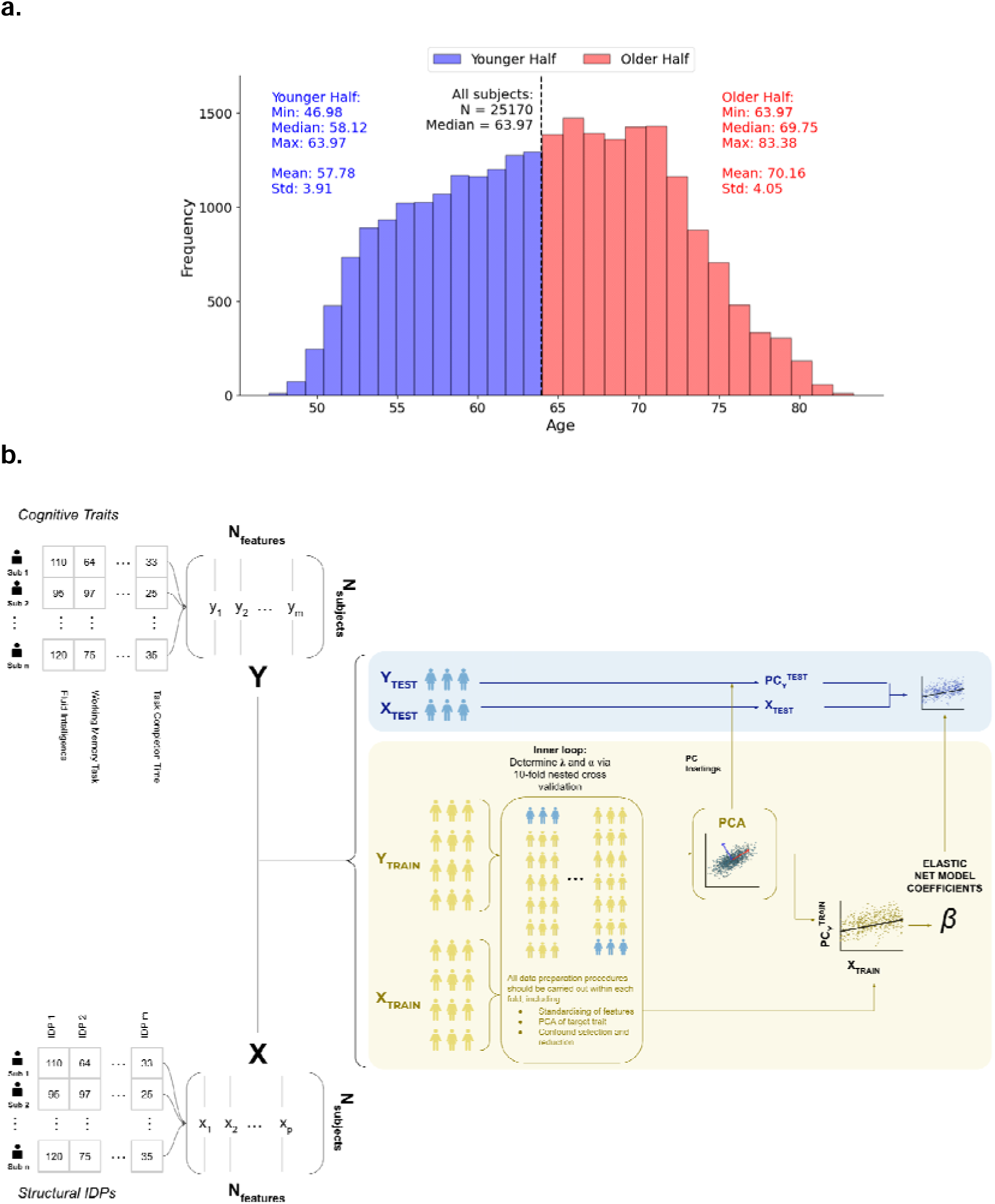
**(a)** Age distribution of UKB subjects, divided into younger (blue) and older (red) halves, each containing 12,585 subjects. Summary statistics indicate that the younger and older halves have nearly identical standard deviations and very similar means relative to their medians, demonstrating a balanced age distribution. **(b)** Cross-validation framework for extracting a composite cognitive measure via PCA while avoiding data leakage. Each subject was represented by a set of structural IDPs (X) and corresponding cognitive trait scores (Y). A nested 10-fold cross-validation framework was used to ensure that all data processing and modelling steps were carried out in a strictly out-of-sample manner. Within each outer fold, the training data underwent an inner loop including standardisation, deconfounding, and elastic net hyperparameter tuning. PCA was applied to the training cognitive scores (Y_TRAIN_), and the first principal component (PC_Y_^TRAIN^) was used as the regression target to determine elastic net coefficients. Separately, the learned PCA loadings were applied to the test data (YTEST) to generate a composite cognitive score (PC_Y_^TEST^), which was used as the out-of-sample target for evaluating model performance using test data IDPs (XTEST) as features.

#### 2.1.1 Confound removal

Large-scale neuroimaging datasets such as UKB have significantly increased the statistical power for detecting small brain-behaviour relationships but have also increased the risk of confounding biases (e.g., age, head motion), which may bias findings and complicate interpretation (Smith & Nichols, 2018). To control for confounding effects, we linearly regressed out the confounds from both imaging and non-imaging variables. Following best practice, we applied deconfounding within cross-validation folds to avoid information from the test set leaking into the training data (Snoek et al., 2019).

We created a reduced set of confounds from a comprehensive set of 559 UKB imaging confounds provided by Alfaro-Almagro et al. (2021; see Supplementary Table SI-2). First, we selected a core subset of conventional confounds including age, sex, age × sex, scanning site, head size, and head motion (see **Supplementary Table SI-2** for the full list), before summarising the remaining confounds (which included additional age-related terms such as age²) using Principal Component Analysis (PCA). Rather than including all 559 variables directly, which would introduce redundancy and multicollinearity, PCA allowed us to capture the shared variance across correlated confounds in a smaller number of orthogonal components. We retained the principal components that together explained greater than 85% of the total variance, consistent with previous work using UKB confounds (Farahibozorg et al., 2021). This ensured that the reduced confound set captured most of the variability of the full set in a lower-dimensional form. In practice, this approach ensured that the influence of the original confounds was still accounted for, while avoiding instability or overfitting that could arise from including hundreds of highly correlated variables directly.

#### 2.1.2 Composite cognitive measure

Prediction accuracy for individual cognitive traits is often limited (Pervaiz et al., 2020), and dividing the sample into age groups further reduces statistical power, limiting our ability to distinguish age-specific effects. Therefore, we developed a composite cognitive measure from UKB traits to improve prediction.

To create a robust cognitive target, we first excluded traits with insufficient data (>50% missing responses). We then ran preliminary elastic net regressions, using all 1,439 IDPs to predict each cognitive measure across all subjects, and ranked the remaining traits by predictive accuracy. As shown in **Supplementary Figure SI-3a** and **SI-3b**, performance dropped sharply beyond the 30th-best trait, motivating the selection of the top 30 traits as candidate targets. The list of these traits is provided in **Supplementary Table SI-3**. Since generalisability can only be meaningfully assessed for traits that are at least moderately predictable, focusing on well-predicted traits ensured that model comparisons reflected meaningful cognitive variation. To maintain consistency across models and avoid the considerable computational cost of evaluating all 1,331 traits in a nested framework, trait selection was performed once, outside the main cross-validated modelling framework.

We then asked whether these top 30 traits captured redundant information or introduced useful diversity. **Supplementary Figure SI-3c** shows that while some traits clustered, many were only weakly related, indicating that the set represents multiple dimensions of cognition rather than a single narrow construct.

To capture the shared variance across these diverse traits and further boost prediction accuracy, we derived a composite by applying PCA, using the first principal component as the target. At this stage, all steps were performed using a fully cross-validated procedure to prevent data leakage (**Figure 1b**). The PCA composite both captured shared variance and achieved higher accuracy than any single trait (**Supplementary Figure SI-3d**), an important property given that prediction accuracy typically decreases after deconfounding.

Importantly, this composite differs from the classical ‘g-factor’: whereas *g* is derived from factor analysis across all available cognitive traits to represent general intelligence, our composite was deliberately restricted to those traits most strongly associated with brain imaging. This composite measure was then used as the target variable for evaluating prediction performance across regression models.

### 2.2 Univariate modelling: age-moderated effects

To investigate how brain-cognition relationships vary across the lifespan, we first applied univariate linear models to test whether the effect of individual IDPs on cognition differs across age quartiles. This section outlines the modelling approach and describes the permutation testing framework used to assess the statistical significance of the effects.

#### 2.2.1 Linear modelling of age-moderated brain-cognition relationships

To estimate age-specific associations between brain structure and cognitive performance, we fitted a linear regression model using as regressors age and a single IDP at a time, treating age as a moderator by stratifying subjects into age quartiles. We created four binary variables representing the age quartiles, assigning a value of 1 for subjects within the respective quartile and 0 otherwise. Each binary variable was multiplied by the selected IDP, creating four new features representing the IDP stratified by age quartile. Each resulting quartile-specific feature was z-scored within the training folds, with the same scaling applied to the test data, ensuring comparability across features and preventing information leakage. Additionally, age was included as a regressor to account for its confounding effect, as is commonly done in neuroimaging studies (Jenkinson & Chappell, 2017; Rao et al., 2015).

#### 2.2.2 Permutation testing of age-moderated effects

To assess statistical differences in the regression coefficients across age quartiles, we used permutation testing. Specifically, for each IDP we fitted a linear regression model across all subjects. This model yielded four quartile-specific IDP coefficients (one for each age quartile), along with an intercept and a continuous age term. We then computed the difference between the coefficient for each quartile and the mean of the other three quartiles.

This procedure was repeated 10,000 times, with subjects’ age-quartile assignments randomly permuted in each iteration, thereby shuffling the mapping between individuals and quartile groups. This generated 10,000 sets of four coefficients per IDP. For each permutation and coefficient, we recalculated the difference between the permuted coefficient and the mean of the remaining three. After all permutations, we computed p-values by determining the proportion of times the permuted difference exceeded the observed difference for each coefficient. This resulted in four p-values for each IDP, one for each quartile.

Under the null hypothesis of no significant coefficient differences across quartiles, the distribution of p-values should be uniform. To assess this hypothesis, we applied a Kolmogorov-Smirnov (KS) test. The KS statistic quantifies the deviation between the empirical cumulative distribution function of the observed p-values and the cumulative distribution function of a uniform distribution expected under the null. A significant KS test for a given age quartile indicates that, across the 1,439 IDPs, the distribution of p-values deviates from uniform, consistent with systematic differences in IDP regression coefficients and therefore age-related variation in brain-cognition relationships. To account for the large number of tests across 1,439 IDPs and four quartiles, we applied Benjamini-Hochberg false discovery rate (FDR) correction to the resulting p-values. This provided a set of corrected significance values for each IDP-by-quartile interaction effect.

### 2.3 Multivariate Modelling: Elastic Net Prediction

We then turned to multivariate prediction, using elastic net regression to predict cognition from all 1,439 structural IDPs simultaneously. Unlike the univariate analyses in **Subsection 2.2**, which examined IDPs one at a time, this approach incorporates all structural features together, allowing the model to capture brain-wide patterns associated with cognition and improving predictive performance in a way that aligns more closely with real-world or clinical applications. Here, we describe the prediction framework used to assess model generalisation across different age groups.

#### 2.3.1 Multivariate prediction using elastic net regression

Elastic net is a common approach for handling multicollinearity in large imaging datasets while avoiding overfitting. Elastic net combines the L_1_ penalty of lasso (Tibshirani, 1996) and the L_2_ penalty of ridge regression (Hoerl & Kennard, 1970), enabling both coefficient shrinkage and sparsity in the model (Vidaurre et al., 2013; Zou & Hastie, 2005).

Let *X* ϵ ℝ^*n×p*^ be the design matrix with *n* subjects and *p* features, and let *y* ϵ ℝ*^n^* be the target vector, representing the composite cognitive measure. Following the implementation in `scikit-learn`, the elastic net regression coefficients, *β* are determined by:

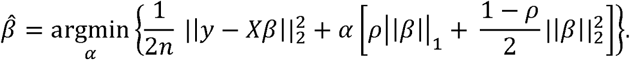

Here, *α* is the overall regularisation strength, and *p* equivalent to `l1_ratio` in scikit-learn) controls the mix between L1 and L2 penalties. Both hyperparameters were selected via nested cross-validation, and all IDPs were z-scored within each training fold (scaling applied to test folds) to ensure comparability across predictors.

#### 2.3.2 Age-split training and generalisation performance

Using a fully cross-validated framework, we applied elastic net regression with 1,439 IDPs to predict cognition, dividing the population into two equally sized age groups (younger and older halves). We opted for two groups, rather than four as in the univariate analyses, to simplify the analysis and ensure larger sample sizes per group. This grouping improved model stability and statistical power during cross-validation, while still capturing meaningful age differences. Prior to modelling, all confounds (including age and age-related terms such as age²) were regressed out from both predictors and the cognitive target (s**ee Subsection 2.1.1**), ensuring that predictions reflected brain-cognition associations independent of known confound effects, consistent with recommended best practice in large-scale neuroimaging analyses. Model performance was evaluated using 10-fold cross-validation, repeated 10 times with different random partitions of the data, resulting in 100 independent estimates of out-of-sample performance.

The population was divided into two equally sized age groups, a younger half and an older half. We examined six distinct training and testing approaches, as shown in **Figure 2**:

1. **Within-age-group models:** Training and testing within the same age group

a. Train: Young; Test: Young
b. Train: Old; Test: Old
2. **Across-age-group models:** Training in one age group and testing in the other

a. Train: Young; Test: Old
b. Train: Old; Test: Young
3. **Pooled models:** Training on the entire dataset and testing within each specific age group

a. Train: ALL; Test: Young
b. Train: ALL; Test: Old

**Figure 2.**
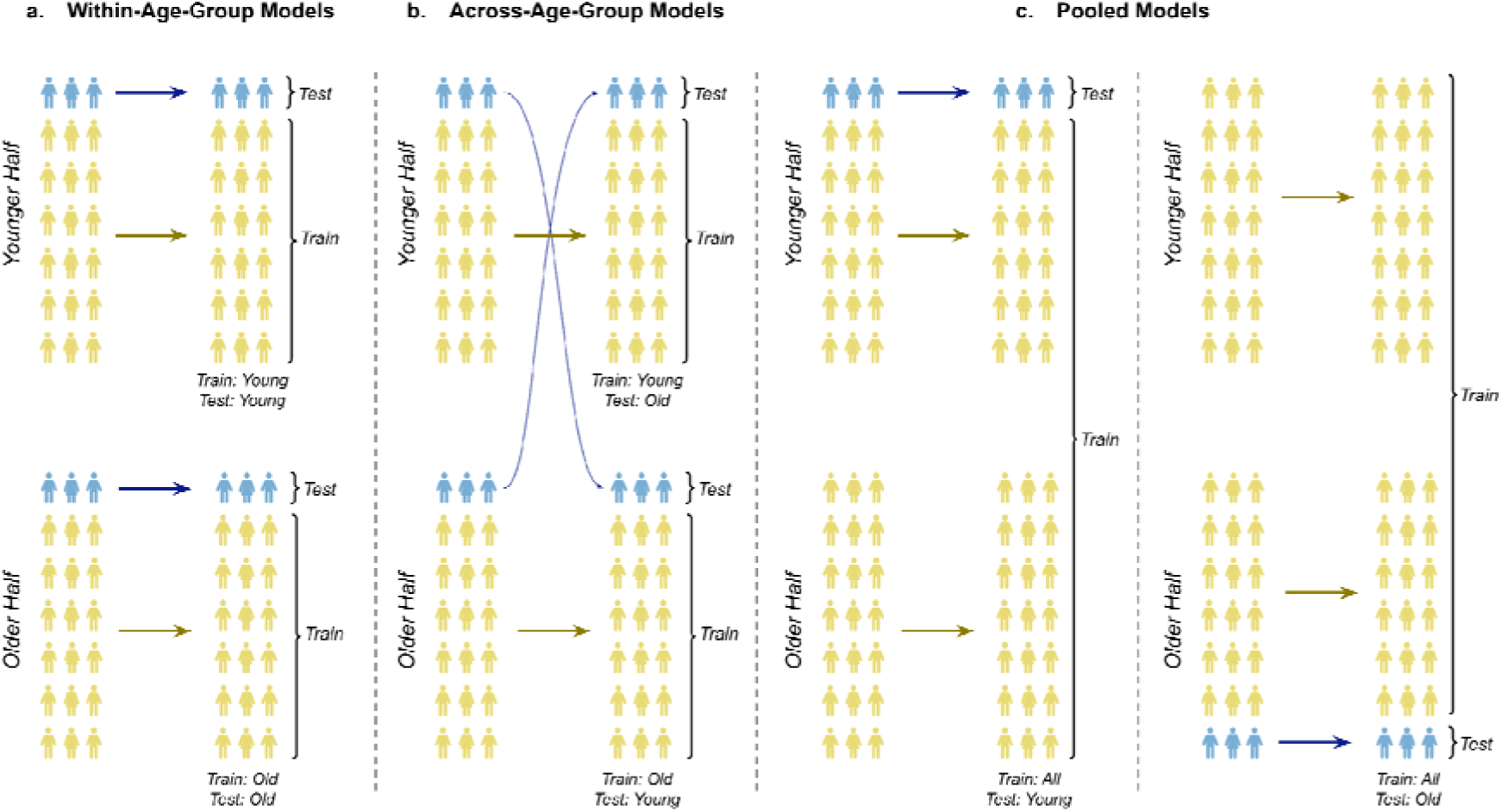
Example fold within the cross-validation framework for predicting cognition from structural IDPs using different model approaches. Three approaches were implemented: **(a) Within-age-group models:** Elastic net regression was applied within the same age group for both training and testing (e.g., Train: Young; Test: Young). **(b) Across-age-group models:** Models were trained on one age group and tested on the other (e.g., Train: Young; Test: Old). **(c)** Pooled models: Models were trained on subjects spanning the full age range, randomly subsampled to match the training sample sizes of the other approaches, with testing restricted to a specific age group (e.g., Train: ALL, Test: Old). This schematic presents a single fold of the 10-fold cross-validation procedure. Each model was trained and tested on an equal number of subjects per fold across all approaches.

To ensure comparability across models, we matched the sample size of the Train: ALL dataset to each age-specific training set by randomly subsampling subjects. This provided a standardised framework for evaluating out-of-sample predictions across all approaches.

#### 2.3.3 Evaluation Metrics

We report Pearson correlation (r), widely used in neuroimaging prediction studies (Vieira et al., 2022) as the primary accuracy metric, because it is invariant to differences in mean and variance and therefore suitable for assessing generalisability across age groups. In contrast, RMSE is sensitive to absolute group differences, and while expressing performance as percent improvement over a null predictor (%ΔRMSE) adjusts for variance, it remains affected by mean shifts between groups. %ΔRMSE is therefore best viewed as a complementary measure, not a replacement for correlation:

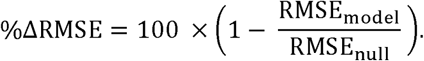

Here, RMSE_null_ equals the standard deviation of the test target, i.e., the variability in cognition that a naïve mean predictor cannot reduce. Given the modest effect sizes typical in brain-cognition prediction (Cox et al., 2019; Rasero et al., 2021; Schulz et al., 2024), absolute improvements in %ΔRMSE are expected to be small. Note that absolute accuracies are further reduced in our design because splitting the cohort into younger and older halves reduces the training sample size by half, but this is necessary to evaluate age-dependent generalisability. For these reasons, we present correlation as the headline metric in the main text, while RMSE-based metrics are provided in the Supplementary Information for completeness.

## 3 Results

We assessed how age moderates the relationship between brain structure and cognition by (i) testing associations between individual IDPs and cognition across age quartiles, and (ii) evaluating the generalisability of multivariate predictive models between age groups.

### 3.1 Age moderates brain-cognition associations

Using the quartile-stratified regression framework described in Methods, **Subsection 2.2.1**, we compared IDP-cognition associations across age groups. This analysis revealed distinct, age-dependent patterns in the regression coefficients linking brain structure to cognition. As shown in **Figure 3a**, certain IDPs exhibited distinct age-related patterns: for example, cortical thickness IDPs tend to show stronger positive coefficients in the youngest quartile (indicated in red) and stronger negative coefficients in the oldest quartile (indicated by blue). **Figure 3b** presents scatterplots comparing coefficients between quartiles, highlighting the particularly low Pearson correlation between the youngest (Q1) and oldest (Q4) quartiles (*r* = 0.40). Importantly, the variance of the cognitive outcome was highly similar across quartiles (SD ≈ 2.1-2.3; **Supplementary Table SI-4**, **Supplementary Figure SI-4**), and IDPs were standardised within each quartile, confirming that coefficient differences are not driven by unequal scaling. These findings are further supported by **Supplementary Figure SI-5a**, which shows a violin plot summarising the full coefficient distributions across quartiles, illustrating both the systematic shift in medians and the increased variability in Q4.

**Figure 3.**
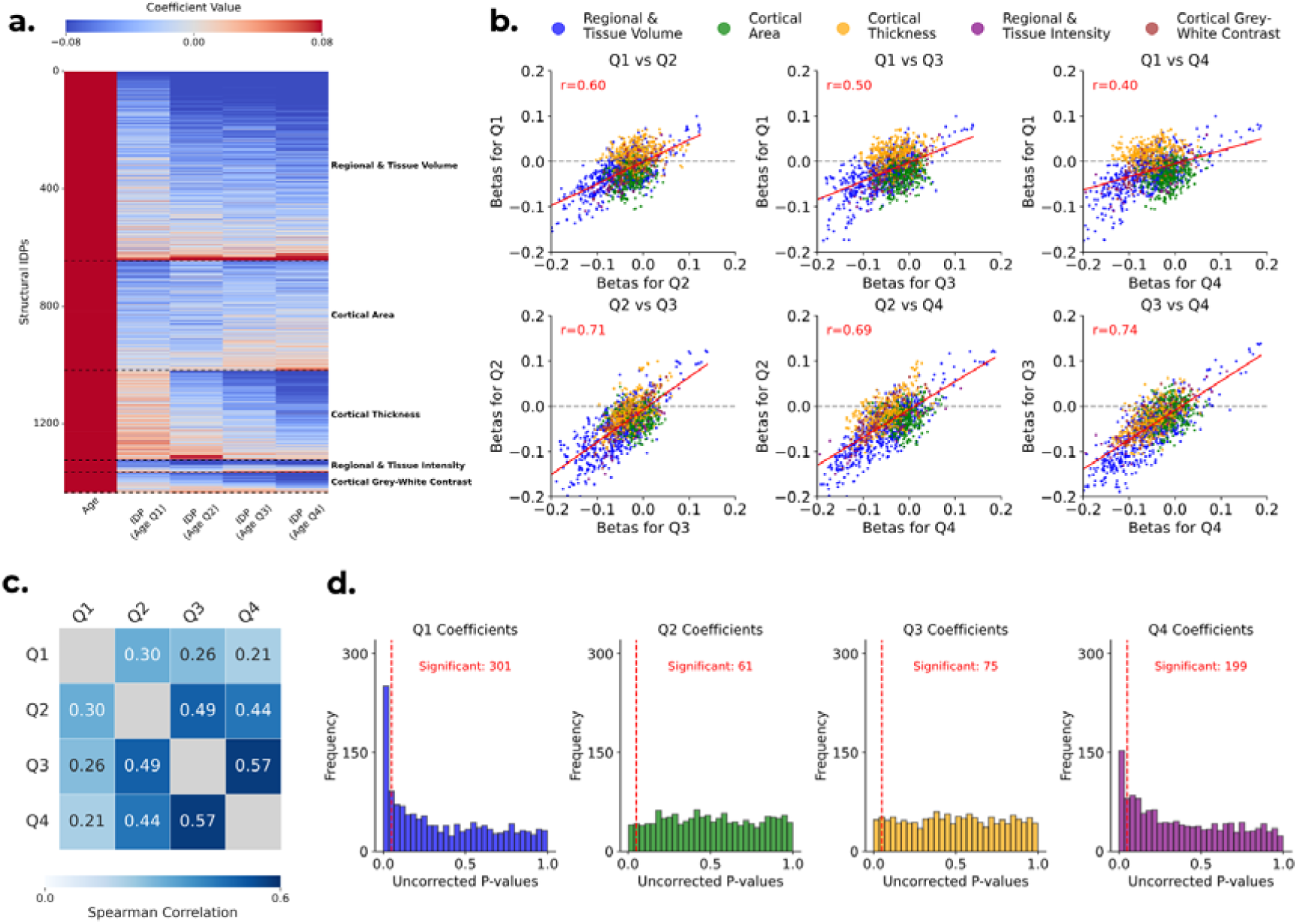
**(a)** Heatmap of coefficients from the linear regression model predicting cognitive performance using each of the 1,439 IDPs individually. The model estimated one coefficient for age and four quartile-specific IDP coefficients (x-axis), shown across all 1,439 IDPs (y-axis). IDPs are categorised into five groups: Regional and Tissue Volume, Cortical Area, Cortical Thickness, Regional and Tissue Intensity, and Cortical Grey-White Contrast. **(b)** Scatterplots comparing regression coefficients for IDPs between age quartiles. Each scatterplot compares coefficients between two quartiles: (i) Q1 vs. Q2. (ii) Q1 vs. Q3. (iii) Q1 vs. Q4. (iv) Q2 vs. Q3. (v) Q2 vs. Q4. (vi) Q3 vs. Q4. **(c)** Heatmap of Spearman’s correlations between the absolute values of all IDP coefficients across age quartiles. Darker shades indicate stronger positive correlations, highlighting the particularly low correlation between the youngest quartile (Q1) and the others. **(d)** Histograms of uncorrected p-values from permutation testing of the IDP–cognition regression coefficients, with one histogram per age quartile (each based on 1,439 IDPs, i.e. one p-value per IDP in that quartile). The red dashed lines mark the p = 0.05 threshold. Under the null hypothesis of no coefficient differences across age quartiles, each distribution would be uniform, and approximately 72 p-values (5% of 1,439) would fall below this threshold by chance.

The spread of scatter points suggested that regression coefficients varied in both size and direction across quartiles. Beyond the direction of these effects, we next asked whether the relative importance of different brain features changed with age. To investigate this, we used the absolute value of the coefficients as a proxy for feature importance and computed the Spearman’s correlation between quartiles. **Figure 3c** shows the resulting heatmap, indicating that the most predictive IDPs in Q1 (i.e., the youngest subjects) differ from those in older subjects. Specifically, the correlations between Q1 and the other quartiles were particularly low (Q1 vs. Q2: r_s_ = 0.30; Q1 vs. Q3: r_s_ = 0.26; Q1 vs. Q4: r_s_ = 0.21), further emphasising the weak resemblance between Q1 and the other age groups. The corresponding Pearson analysis is shown in **Supplementary Figure SI-5b**, which also highlights the differences between coefficients of Q1 and the remaining quartiles.

Permutation testing of the IDP-cognition regression coefficients (see **Subsection 2.2.2**) produced 5,756 p-values (1,439 IDPs × 4 quartiles). **Figure 3d** shows the distributions, which deviated from uniformity for the youngest and oldest quartiles (Q1: *p*_KS_ < 0.001; Q4: *p*_KS_ < 0.001), but not for the middle quartiles (Q2: *p*_KS_ = 0.20; Q3: *p*_KS_ = 0.73). This indicates that age-moderated effects were concentrated at the extremes of the age distribution.

Having established that age moderates the brain-cognition relationship for at least some of the IDPs (particularly in the youngest and oldest quartiles), we next identified which IDPs remained significant after FDR (see **Subsection 2.2.2**). The corrected p-value distributions are shown in **Supplementary Figure SI-5c**. Although the number of significant IDPs was modest after correction, a subset remained significant in the youngest and oldest quartiles, with 26 IDPs in Q1 and 13 in Q4, with some appearing in both. In total, 35 unique IDPs were significant, falling into the following categories: Cortical Thickness (26), Regional and Tissue Volume (7), and Regional and Tissue Intensity (2). A full list of significant IDPs is provided in **Supplementary Table SI-5**. Consistent with chance expectations under the null, the middle quartiles showed no significant effects after correction, aside from one regional and tissue volume IDP in Q3. This emphasises that age-moderation was strongest at the extremes of the age distribution.

In summary, these findings reveal that, beyond the general effects of age, the relationship between structural IDPs and cognitive performance varies across age groups. These results therefore support our hypothesis that between-subject variations in brain structure associated with variations in cognition in older people may be different from those in younger people. **Supplementary Figure SI-5d** further illustrates this by showing Bland-Altman plots of Fisher-transformed IDP-cognition correlations across age quartiles, revealing systematic shifts and widening differences in association strength between Q1 and Q4.

### 3.2 Age-specific models outperform pooled and cross-age generalisation

Using the multivariate elastic net framework described in **Subsection 2.3**, we systematically compared the out-of-sample prediction accuracy of the three training strategies: within-age-group, across-age-group, and pooled models. This analysis revealed a clear and consistent performance hierarchy, as shown in **Figure 4a**. Age-specific models achieved the highest accuracy (r_WITHIN-AGE-GROUP_ = 0.095), significantly outperforming both pooled models (r_POOLED_ = 0.090; p = 0.002) and, most strongly, models trained on a different age group (r_ACROSS-AGE-GROUP_ = 0.085; p < 0.001). Although these effect sizes are modest, they were consistent and statistically reliable. Pooled models, trained on a mix of ages, offered a robust intermediate performance, in turn proving more accurate than the across-age-group models (p = 0.02). This consistent ordering (**within-age-group > pooled > across-age-group**) highlights the clear advantage of training models on data that is most representative of the test population.

**Figure 4.**
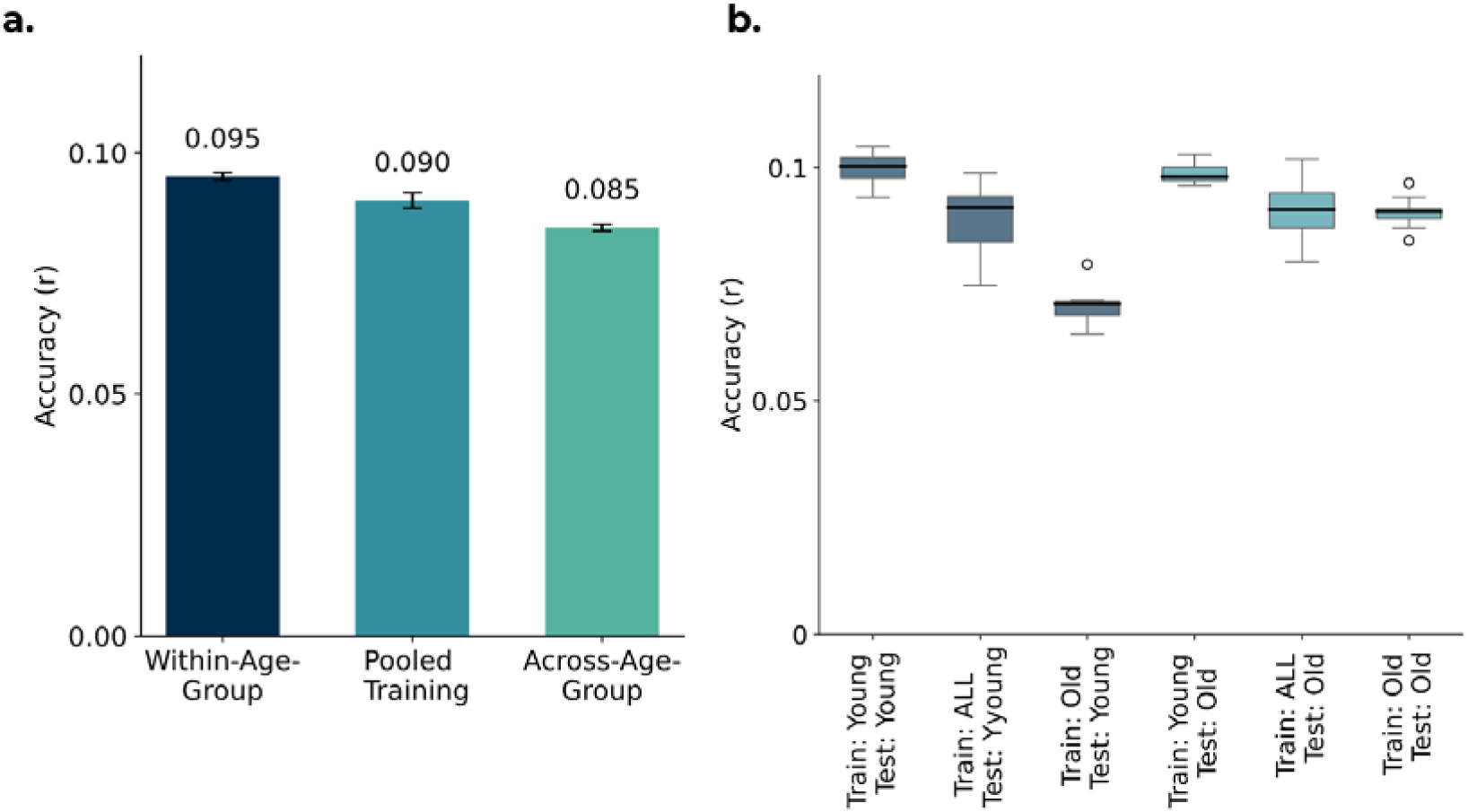
Comparison of model performance, regularisation strength, and regression coefficients across different training strategies when using 1,439 structural IDPs to predict cognition, evaluated using 10-fold cross-validation repeated 10 times (100 out-of-sample estimates). Both the IDPs and cognitive trait were deconfounded for all confounds (including age-related confounds). Accuracy is measured as the Pearson correlation (r) between predicted and observed cognitive scores. **(a)** Bar chart comparing prediction accuracy for within-age-group training and testing (Train: Young, Test: Young; and Train: Old, Test: Old), across-age-group training and testing (Train: Young, Test: Old and Train: Old, Test: Young), and pooled training (Train: ALL, Test: Young; and Train: ALL, Test: Old). Error bars indicate the standard deviation across the 10 repeats. * p < 0.05; ** p < 0.01; *** p < 0.001. **(b)** Comparison of prediction accuracy across six training/testing splits, with models trained and tested on different age groups. Boxplots show the distribution across cross-validation repetitions.

Consistent patterns were observed when accuracy was expressed as improvement over a null RMSE. Within-age-group models achieved the highest gains, followed closely by pooled models, while across-age-group models performed substantially worse. Absolute improvements were small (<1%) as expected given the modest effect sizes typical in this domain (**Supplementary Figure SI-6**).

### 3.3 Younger-trained models generalise across age groups

We next examined whether the observed effects were symmetrical across age groups. **Figure 4b** shows that models trained on younger subjects consistently outperformed those trained on older subjects, regardless of the testing group. Although the gap was modest when predicting older subjects (*r*_TR: Y; TE: O_ = 0.099 vs. *r*_TR: O; TE: O_ = 0.090; p < 0.001), the gap was more pronounced for younger subjects, where older-trained models were notably less accurate (*r*_TR: Y; TE: Y_ = 0.100 vs. *r*_TR: O; TE: Y_ = 0.070; p < 0.001). This suggests that models trained on older subjects struggle to capture brain-cognition relationships relevant to younger individuals, whereas those trained on younger subjects remain predictive even for older individuals.

**Figure 4bError! Reference source not found.** further shows that the effect of training strategy depended on the test group. For younger subjects, within-age-group models achieved the highest accuracy, outperforming both pooled (r_TR: Y; TE: Y_ = 0.100 vs. r_TR: ALL; TE: Y_ = 0.089; p < 0.001) and across-age-group models (r_TR: Y; TE: Y_ = 0.100 vs. r_TR: O; TE: Y_ = 0.070; p < 0.001). For older subjects, performance was more similar across conditions: within-age-group and pooled models performed comparably (r_TR: O; TE: O_ = 0.090 vs. r_TR: ALL; TE: O_ = 0.091; p = 1.00), while younger-trained models outperformed pooled models (r_TR: Y; TE: O_ = 0.099 vs. r_TR: ALL; TE: O_ = 0.091; p = 0.006).

RMSE-based accuracy metrics are provided in **Supplementary Figures SI-6c** and **SI-6d**. These broadly supported the correlation-based findings, while also illustrating how between-group variance differences can affect RMSE-based measures.

Overall, these results demonstrate a key asymmetry in generalisation, where brain-cognition patterns from a younger population serve as a more generalisable model, while patterns unique to an older population do not. These analyses highlight a trade-off between model generalisability and age-group specificity.

### 3.4 Older-trained models need to compensate higher noise

To understand the source of these performance differences, we examined the models’ hyperparameters. As shown in **Figure 5a**, models trained on older subjects consistently selected higher levels of regularisation (α) than those trained on the full population, both when tested on younger subjects (α_TR: O; TE: Y_ = 7.87 vs. α_TR: ALL; TE: Y_ = 4.97) and on older subjects (α_TR: O; TE: O_ = 8.40 vs. α_TR: ALL; TE: O_ = 5.17). By contrast, models trained on younger subjects used lower levels of regularisation, closer to pooled models (α_TR: Y; TE: O_ = 5.00; α_TR: Y; TE: Y_ = 6.20).

**Figure 5.**
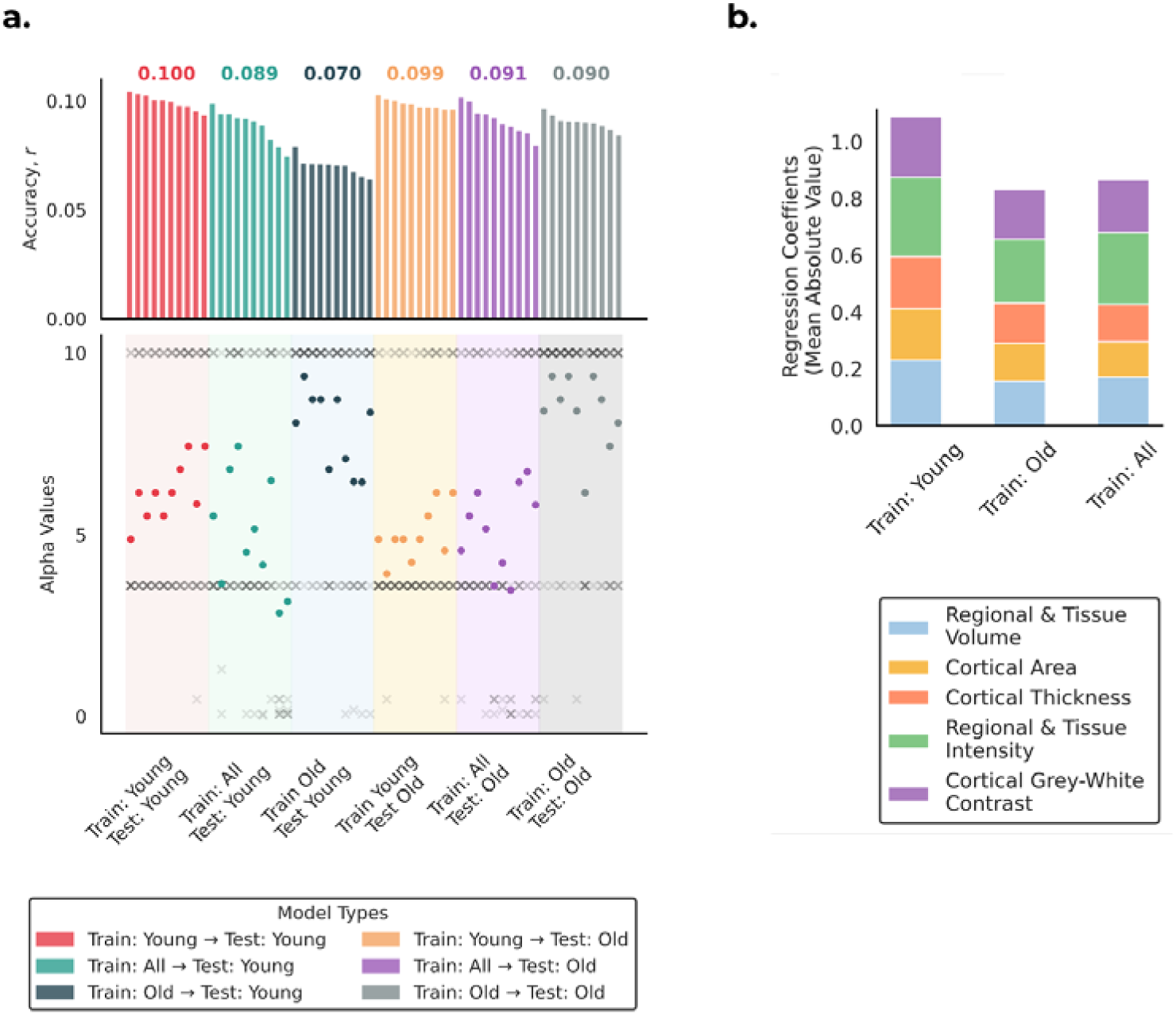
Overview of model regularisation and feature importance across training strategies. **(a)** Comparison of elastic net regularisation strength and model accuracy. The top panel shows mean prediction accuracy across cross-validation repetitions (numbers above bars indicate the mean r). The bottom panel shows the regularisation strength (α) per cross-validation repetition (× = individual folds; ● = mean across 10 folds). Hyperparameters were determined via nested cross-validation, with the inner loop selecting the optimal α. **(b)** Stacked bar plot showing the mean absolute values of regression coefficients, averaged across IDPs within each structural category, for the three training approaches.

These results suggest that models trained on older subjects tend to select stronger regularisation to avoid overfitting, consistent with higher noise levels or greater inter-subject variability in this group. Models trained on younger subjects, or on pooled samples that include younger individuals, require less regularisation, which may explain why pooled models perform comparably to those trained only on older subjects. Although including younger subjects in pooled models may introduce distributional bias when testing on older subjects, their lower noise levels reduce the need for regularisation and help preserve predictive signal.

An analogous analysis of the elastic-net mixing parameter (ρ; equivalent to the L1 ratio in scikit-learn) is provided in **Supplementary Figure SI-7**. Age-specific models (Train: Young and Train: Old) consistently selected ridge-like solutions (L1 ratio ≈ 0, i.e. favouring shrinkage of all coefficients rather than sparsity), whereas pooled models showed small but non-zero L1 ratios (0 to 0.25). This indicates that the principal difference across training strategies lies in the strength of regularisation (α), rather than in the balance between L1 and L2 penalties.

**Figure 5b** shows that models trained on older subjects had consistently smaller mean absolute regression coefficients across IDP categories, relative to younger-trained models. The largest relative reductions were observed in Regional and Tissue Volume (31.7% across 646 IDPs), Cortical Area (26.5% across 372 IDPs), and Cortical Thickness (22.9% across 306 IDPs). Smaller but consistent reductions appeared in Regional and Tissue Intensity (19.8% across 41 IDPs) and Cortical Grey-White Contrast (16.9% across 70 IDPs). Given the larger number of IDPs in the first three categories, these reductions are likely to have the greatest impact on model performance. Pooled training produced intermediate values across all categories, consistent with the idea that including a broader age range stabilises coefficient estimates by averaging over age-specific variability.

## 4 Discussion

Accounting for the influence of age is essential when studying brain-cognition relationships. While the standard approach is to remove age as a confound, we suggest it is more appropriate to treat age as a moderator, finding that the statistical links between brain structure and cognition vary significantly across the lifespan. Although absolute prediction accuracies were modest (r ≈ 0.07-0.10), they are consistent with prior large-sample neuroimaging studies after accounting for sample size and deconfounding (Cox et al., 2019; Rasero et al., 2021; Schulz et al., 2024). Our focus was not maximising absolute accuracy, but assessing whether the brain-cognition relationships generalise across age groups. This led to an important finding: an asymmetry in how predictive models generalise across age.

### Generalisation asymmetry and its link to age-related variability

A central finding is an asymmetry in how predictive models generalise across age. Models trained on younger subjects generalised well to older subjects, likely capturing foundational brain-cognition relationships that remain stable across the lifespan. In contrast, models trained on older subjects generalised poorly to younger individuals.

Our analysis of the model’s hyperparameters provides an explanation for this failure. Older-trained models consistently selected stronger regularisation to prevent overfitting, which is consistent with higher noise levels (Nomi et al., 2024) or greater inter-subject variability in this group. Including younger subjects, as in the pooled models, appears to stabilise coefficients by reducing this need for high regularisation. This suggests that the brain-cognition relationship in older age is complicated by additional sources of variance—such as differential neurodegeneration or comorbidities—making it harder to learn a pattern that generalises back to the more homogeneous younger population. Univariate analyses also supported these findings: a subset of IDPs showed significant coefficient differences across quartiles, and within-age-group models performed better than across-age-group models. This highlights the benefit of training on data matched to the test set distribution.

Previous studies have reported age-related differences in prediction generalisability (Yu & Fischer, 2022); we extend these effects using UKB and treat age as a moderator rather than purely as a confound. One approach to model moderation is using age × IDP interaction terms (Burzynska et al., 2012; de Chastelaine et al., 2019, 2023), but this can conflate underlying nonlinearity with interaction effects (Rimpler et al., 2024). We therefore adopted an age-stratified predictive framework that tests whether associations learned in one age group generalise to another.

### Training strategy

While within-age-group models achieved the highest accuracy when tested on their own age group, they generalised poorly to the other group. In practice, age-specific models may be useful for targeted subgroups (e.g., younger adults), but their performance can be inconsistent out-of-distribution, particularly when trained on older adults and tested on younger ones. By contrast, pooled models did not reach the very highest within-group accuracy but provided more consistent predictions across both age groups, consistent with the idea that including younger subjects with more stable brain–cognition relationships reduces variability. For older groups, a more diverse training set improved generalisation without sacrificing within-group accuracy. Overall, pooled training (i.e., the conventional approach) offered the most robust performance for heterogeneous datasets or limited sample sizes, helping to avoid overfitting to a specific age group while maintaining stable performance.

### Limitations and next steps

We concentrated on structural IDPs for their robust age trends, high test-retest reliability, larger UKB sample coverage than fMRI, and reduced sensitivity to age-related motion. However, we acknowledge that functional connectivity measures can yield higher prediction accuracy (Ooi et al., 2022; Soch et al., 2022).

In this work, we focused on the first principal component derived from cognitive traits. However, cognition comprises multiple subtypes (e.g., crystallised and fluid intelligence), which could be examined in addition.

Poorer generalisation in older-trained models may partly reflect the higher prevalence of age-related comorbidities, e.g., hypertension, diabetes, cardiac disease. These are known to affect brain IDPs; restricting to particularly healthy participants or deconfounding such conditions would be a valuable direction for future work. Furthermore, we recognise that the prevalence of comorbidities and other age-related factors varies across populations. Replication in more diverse cohorts will be important to test the broader generalisability of our findings.

Finally, although we divided subjects into halves and quartiles, a continuous age-varying coefficient model could capture nonlinear effects more precisely and help identify specific ages at which brain–cognition relationships change. Combining this with more advanced deconfounding methods, such as propensity score matching (Rosenbaum & Rubin, 1983) or confound-isolating cross-validation (Chyzhyk et al., 2022), may capture nonlinear effects more precisely and improve robustness by reducing residual bias (Power et al., 2012).

### Summary

This work demonstrates that the link between brain structure and cognition is not inalterable but changes across the adult lifespan. A central finding was that there is an asymmetry in generalisability, with models trained on younger subjects able to predict cognition in older individuals, but models trained on older subjects unable to generalise to younger individuals. This reveals an important trade-off for researchers and clinicians building predictive models, with the optimal approach depends directly on the intended goal.

For example, in situations requiring high accuracy within a specific target group (e.g., clinical diagnostics), an age-specific model may be preferable. However, in situations requiring a model to generalise reliably across a diverse, mixed-age population, a pooled model trained on all ages provides more stable and broadly applicable predictions. Recognising this trade-off is essential for improving prediction models in both research and clinical settings, ensuring that predictions are optimal for the intended population.

## Data and Code Availability

The data used in this manuscript pertains to a well-known, open-access repository: the UK Biobank dataset. Analysis code is available from the corresponding author upon reasonable request.

## Author Contributions

B.G.: Conceptualisation, Methodology, Validation, Formal Analysis, Writing — Original Draft, Visualisation. C.G.: Software, Writing — Reviewing & Editing. M.W.W.: Methodology, Supervision, Writing — Reviewing & Editing. S.M.S.: Methodology, Supervision, Writing — Reviewing & Editing. D.V.: Conceptualisation, Methodology, Supervision, Writing — Reviewing & Editing, Funding Acquisition.

## Ethics Statement

Participants were drawn from an open-access dataset, UK Biobank, for which all participants provided informed consent, and the established procedures to access and use the data were followed.

## Declaration of Competing Interest

The authors declare no competing interests.

## Acknowledgements

D. Vidaurre is supported by a Novo Nordisk Foundation Emerging Investigator Fellowship (NNF19OC-0054895), an ERC Starting Grant (ERC-StG-2019-850404), and a DFF Project from the Independent Research Fund of Denmark (2034-00054B).

This research has been conducted in part using the UK Biobank Resource under Application Number 8107. We are grateful to UK Biobank for making the data available, and to all UK Biobank study participants, who generously donated their time to make this resource possible.

Analysis was carried out on the clusters at the Oxford Biomedical Research Computing (BMRC) facility. BMRC is a joint development between the Wellcome Centre for Human Genetics and the Big Data Institute, supported by Health Data Research UK and the NIHR Oxford Biomedical Research Centre.

This research was funded in part by the Wellcome Trust (215573/Z/19/Z). For the purpose of Open Access, the author has applied a CC BY public copyright licence to any Author Accepted Manuscript version arising from this submission.

MWW’s research is supported by the Wellcome Trust (106183/Z/14/Z, 215573/Z/19/Z), the New Therapeutics in Alzheimer’s Diseases (NTAD) study supported by UK MRC, the Dementia Platform UK (RG94383/RG89702) and supported by the NIHR Oxford Health Biomedical Research Centre (NIHR203316). The views expressed are those of the author(s) and not necessarily those of the NIHR or the Department of Health and Social Care. The Wellcome Centre for Integrative Neuroimaging is supported by core funding from the Wellcome Trust (203139/Z/16/Z and 203139/A/16/Z).

## Notes

### Competing Interest Statement

The authors have declared no competing interest.

### Summary of Updates

This version includes revised abstract and introduction text, expanded methods with additional metrics, updated results and discussion, and new supplementary figures.

