## Supplementary Material for "The role of age in the relationship between brain structure and cognition: moderator or confound?"

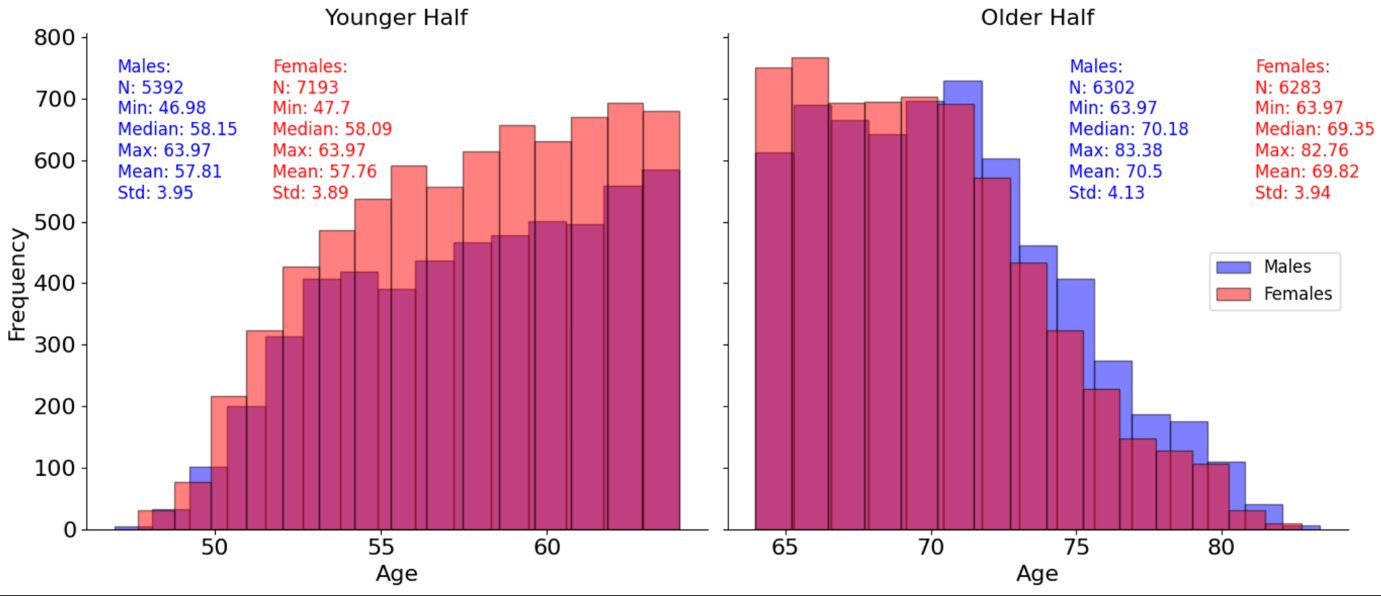

**Figure SI-1.** Histogram of age distributions in UKB subjects, split by sex within each age group. Both younger and older halves contain approximately equal numbers of males and females, with similar age distributions across sexes. This confirms that observed effects are unlikely to be driven by imbalanced age-sex sampling.

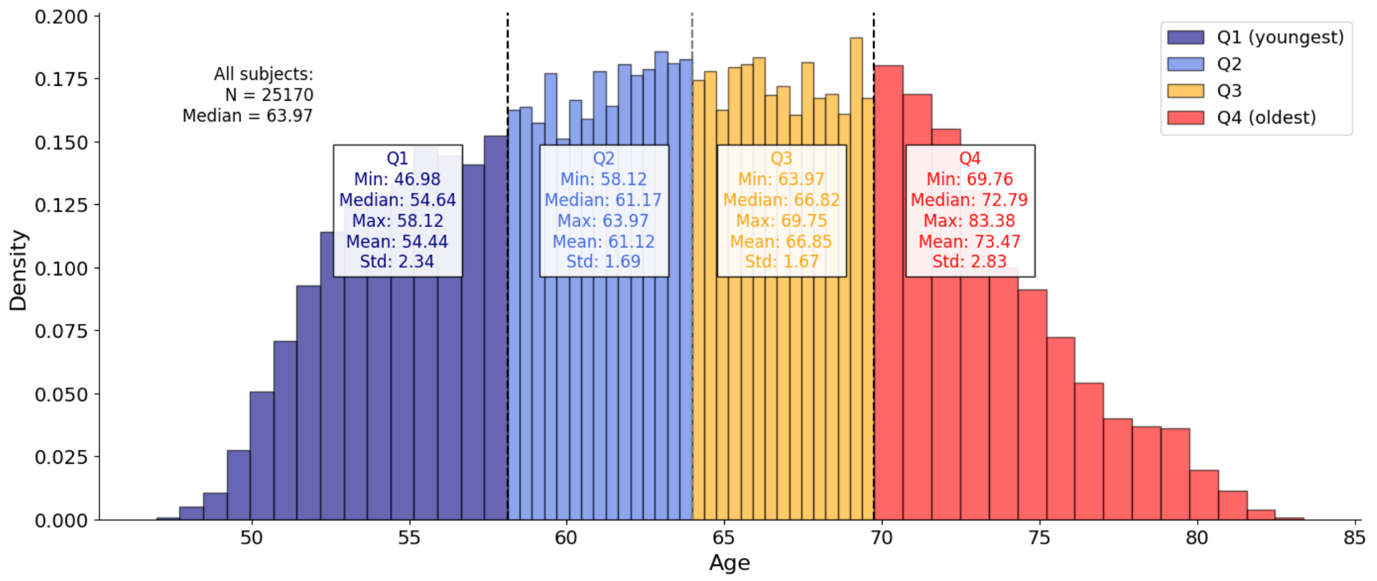

**Figure SI-2.** Histogram of participant age distributions, divided into quartiles. Each colour-coded section corresponds to one of the four age quartiles used in the analysis (Q1: youngest, Q4: oldest). Vertical dashed lines indicate the boundaries between quartiles.

**Table SI-1.** Summary of confound groups used in the analysis, along with the number of variables in each group (n = 559).

| **Confound no.** | **Confounds Group** | **Number of Confounds** |
| --- | --- | --- |
| 1 | Age | 3 |
| 2 | Sex | 3 |
| 3 | Age by Sex | 3 |
| 4 | Head Size | 3 |
| 5 | Site | 2 |
| 6 | Batch | 16 |
| 7 | Protocol | 7 |
| 8 | Service Pack | 1 |
| 9 | Freesurfer T2 | 3 |
| 10 | Scaling | 9 |
| 11 | Echo Time | 6 |
| 12 | Structural Motion | 3 |
| 13 | Table Position | 12 |
| 14 | Nonlinear Registration | 134 |
| 15 | Crossed Terms | 216 |
| 16 | Acquisition Time | 59 |
| 17 | Acquisition Date | 82 |

**Table SI-2.** List of the 14 essential confounds used in the primary deconfounding model.

| **Confound no.** | **Essential Confound Name** |
| --- | --- |
| 1 | Age at Site 1 |
| 2 | Age at Site 2 |
| 3 | Age at Site 3 |
| 4 | Age by Sex at Site 1 |
| 5 | Age by Sex at Site 2 |
| 6 | Age by Sex at Site 3 |
| 7 | Sex at Site 1 |
| 8 | Sex at Site 2 |
| 9 | Sex at Site 3 |
| 10 | Site 1 vs Site 2 |
| 11 | Site 1 vs Site 3 |
| 12 | Head Size at Site 1 |
| 13 | Head Size at Site 2 |
| 14 | Head Size at Site 3 |

To ensure stable and comparable targets, we first excluded cognitive traits with insufficient data, retaining only those with more than 50% valid responses. We then ranked traits by cross-validated brain-cognition correlation and inspected the ranked curve (**Figure SI-3b**), which shows a clear elbow near rank ≈30. Guided by this inflection (and the corresponding correlation cutoff in **Figure SI-3a**), we selected the top 30 traits as candidate targets. The selected traits span a broad range of UKB measures (**Figure SI-3c**), ensuring diversity rather than clustering into a single domain. This ensures that we capture multiple dimensions of cognition rather than reducing to a single narrow trait. Finally, we derived a composite measure by applying PCA to these 30 traits. The first principal component both captures their shared variance and achieves higher prediction accuracy (**Figure SI-3d**). This step was particularly important since prediction accuracy is expected to decrease after deconfounding for age; starting with a strong composite helps ensure the comparisons in the main text are meaningful.

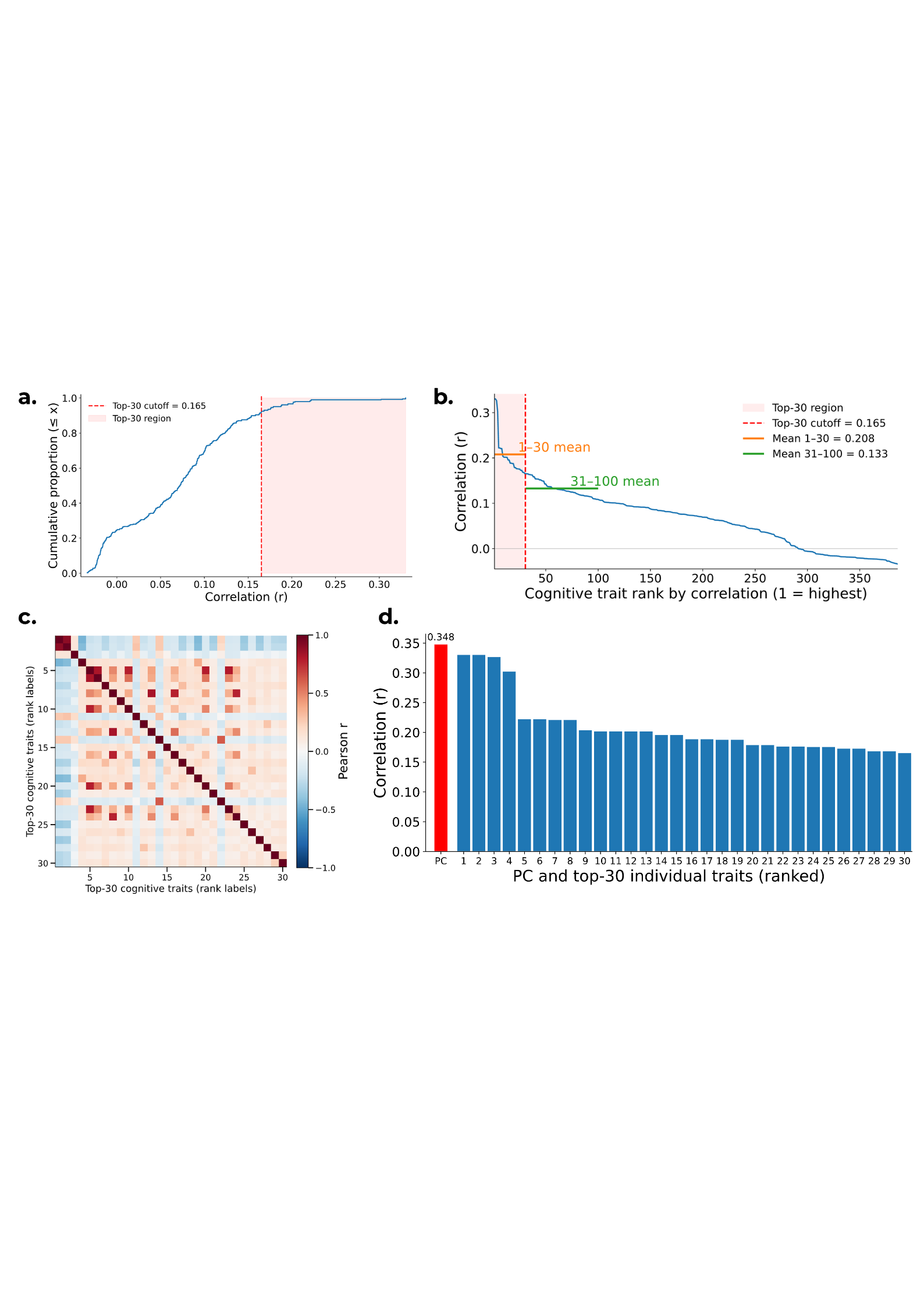

**Figure SI-3.** Selection and reduction of cognitive traits for prediction analyses. **(a)** ECDF of cross-validated brain-cognition correlations for all cognitive traits meeting the coverage criterion (>50% valid responses per trait); the dashed line marks the 30th-highest correlation. **(b)** Ranked plot of individual trait correlations. The dashed line marks rank 30; lines indicate mean correlations for ranks 1-30 and 31-100, showing a clear drop after the top 30. **(c)** Pairwise correlation matrix of the top 30 traits, indicating that while some traits cluster, many are only weakly related, so the set captures diverse cognitive dimensions rather than being redundant. **(d)** Predictive accuracy for the top 30 individual traits (blue) versus the first principal component computed from them (red). The PCA composite captures shared variance and achieves higher accuracy, so we use it as the final cognitive target.

Correlations (rather than error metrics) are shown to ensure comparability across traits with different scales and ranges. Trait selection was performed without deconfounding, first to maximise predictive potential and second to avoid biasing trait selection by confounds that were explicitly tested in later analyses.

**Table SI-3.** Top 30 UK Biobank cognitive traits used to construct the composite cognitive measure, with variable names and corresponding Field IDs.

| **Var. no.** | **Column Header** | **UKB Field ID** |
| --- | --- | --- |
| 1 | Duration spent answering each puzzle (2.0) | 6333 |
| 2 | Duration spent answering each puzzle (2.1) | 6333 |
| 3 | Duration spent answering each puzzle (2.4) | 6333 |
| 4 | Duration spent answering each puzzle (2.7) | 6333 |
| 5 | Duration to complete alphanumeric path (trail #2) (2.0) | 20157 |
| 6 | Duration to complete numeric path (trail #1) (2.0) | 20156 |
| 7 | Duration to entering symbol choice (2.11) | 6325 |
| 8 | Duration to entering symbol choice (2.16) | 6325 |
| 9 | Duration to entering symbol choice (2.20) | 6325 |
| 10 | Duration to entering symbol choice (2.22) | 6325 |
| 11 | Duration to entering symbol choice (2.24) | 6325 |
| 12 | Duration to entering symbol choice (2.4) | 6325 |
| 13 | Duration to entering symbol choice (2.8) | 6325 |
| 14 | Duration to first press of snap-button in each round (0.5) | 404 |
| 15 | Duration to first press of snap-button in each round (0.7) | 404 |
| 16 | Duration to first press of snap-button in each round (2.10) | 404 |
| 17 | Duration to first press of snap-button in each round (2.11) | 404 |
| 18 | Duration to first press of snap-button in each round (2.5) | 404 |
| 19 | Duration to first press of snap-button in each round (2.7) | 404 |
| 20 | Fluid intelligence score (2.0) | 20016 |
| 21 | Mean time to correctly identify matches (0.0) | 20023 |
| 22 | Mean time to correctly identify matches (2.0) | 20023 |
| 23 | Number of puzzles attempted (2.0) | 6374 |
| 24 | Number of puzzles correct (2.0) | 20760 |
| 25 | Number of puzzles correctly solved (2.0) | 6373 |
| 26 | Number of symbol digit matches attempted (2.0) | 23323 |
| 27 | Number of symbol digit matches made correctly (2.0) | 23324 |
| 28 | Number of word pairs correctly associated (2.0) | 20197 |
| 29 | Time to complete round (0.2) | 400 |
| 30 | Time to complete round (2.2) | 400 |

The standard deviations of the cognition score were highly similar across quartiles (≈2.1-2.3). This consistency suggests that differences in regression coefficients across age groups are unlikely to be artefacts of unequal outcome variance, supporting the interpretation that they reflect genuine moderation of brain-cognition associations by age.

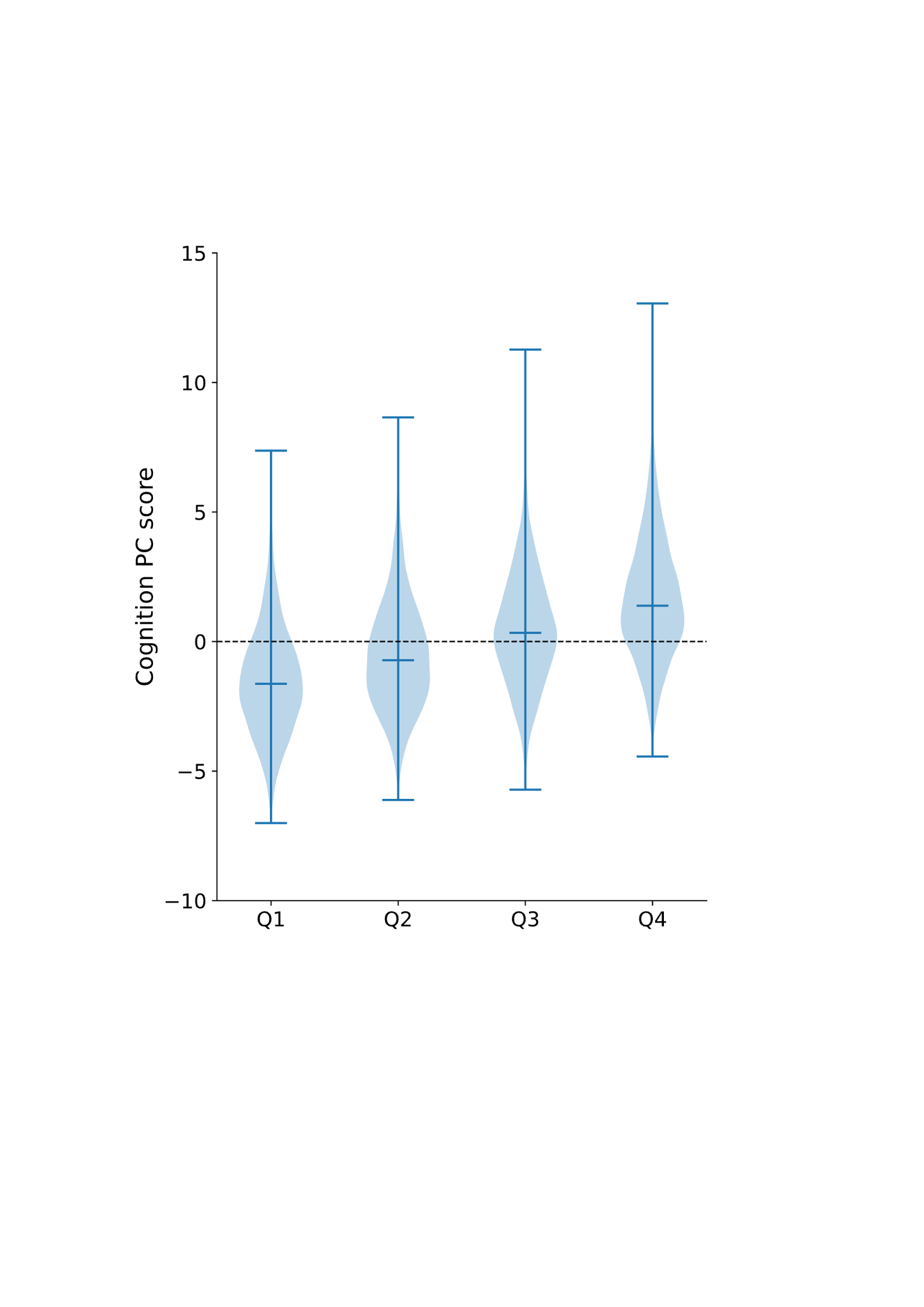

**Figure SI-4.** Violin plots showing the distribution of the composite cognitive measure (PC1 of the top 30 cognitive traits) across age quartiles (Q1-Q4). While mean scores increase systematically with age, the spread of the distributions remains comparable (SD ≈ 2.1-2.3), consistent with Table SI-4. This indicates that observed differences in regression coefficients between quartiles are not artefacts of unequal variance in the cognitive outcome. As with any PCA-derived component, the direction of PC1 is arbitrary, so the systematic shift in means across quartiles reflects relative rather than absolute ordering, and arises from a weighted mix of cognitive traits with different age trends.

**Table SI-4.** Distribution of the composite cognitive measure (i.e., the first PC of the top 30 cognitive traits; PC1) across age quartiles. Shown are sample size (N), mean cognition score, and standard deviation (SD) of cognition scores, along with approximate age ranges for each quartile. While means increase systematically with age, SDs remain similar (≈2.1-2.3), indicating that differences in coefficients between quartiles are not driven by unequal outcome variance.

| **Quartile** | **N** | **Mean** | **SD** | **Age range** |
| --- | --- | --- | --- | --- |
| Q1 | 1957 | -1.51 | 2.09 | ≤ 58.3 |
| Q2 | 1956 | -0.55 | 2.13 | 58.3-64.0 |
| Q3 | 1956 | 0.49 | 2.24 | 64.0-69.7 |
| Q4 | 1956 | 1.64 | 2. 31 | > 69.7 |

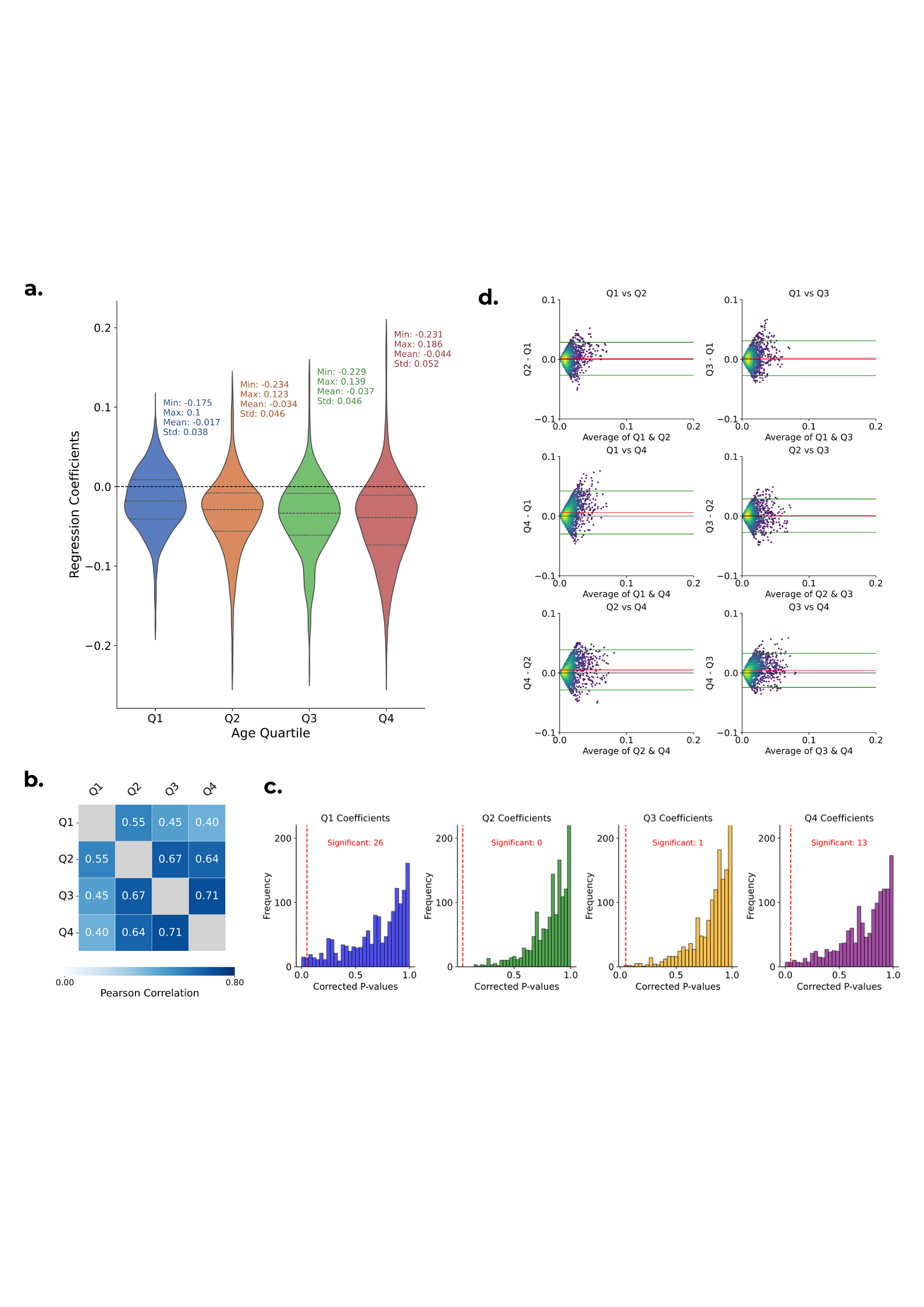

**Figure SI-5.** Summary of how the relationship between structural IDPs and cognition varies across age quartiles. **(a)** Violin plots showing the distribution of ridge regression coefficients across 1,439 IDPs for each age quartile (Q1-Q4), from youngest to oldest. These coefficients are derived from models that include age and the age-split IDP features. The Q4 group exhibits the largest spread and most negative median, suggesting increasing heterogeneity in brain-cognition associations with age. **(b)** Heatmap of Pearson correlations between the absolute values of IDP coefficients across quartiles. Q1 shows particularly low correlation with the other age groups, reinforcing age-specificity in structure-cognition links. **(c)** Histograms of corrected p-values from permutation tests assessing whether each quartile’s coefficient significantly deviates from the mean of the other three. The youngest (Q1) and oldest (Q4) quartiles show the most significant differences after FDR correction (26 and 13 IDPs, respectively), while Q2 and Q3 closely follow the null. **(d)** Bland-Altman plots comparing Fisher-transformed IDP-cognition correlations across pairs of age quartiles. Each plot shows the difference in correlation (y-axis) against the average correlation (x-axis). Mean differences and 95% limits of agreement are shown with dashed lines. Later quartile comparisons (e.g., Q4 vs. Q1) reveal larger divergence in correlation strength.

**Table SI-5.** List of IDPs showing significant age-moderated effects on cognition after FDR correction (Benjamini-Hochberg, q < 0.05), stratified by age quartile. For each quartile, p-values are reported; asterisks indicate significance after FDR correction (* p < 0.05; ** p < 0.01; *** p < 0.001). IDPs are grouped by category.

| IDP Name | Category | Q1 p-val | Q2 p-val | Q3 p-val | Q4 p-val |
| --- | --- | --- | --- | --- | --- |
| IDP_T1_FAST_ROIs_L_cerebellum_crus_II | Regional And Tissue Volume | 0.0261* | 0.6305 | 0.9085 | 0.3633 |
| aseg_rh_volume_VentralDC | Regional And Tissue Volume | 0.0348* | 0.5124 | 0.7701 | 0.8432 |
| HippSubfield_lh_volume_CA1-head | Regional And Tissue Volume | 0.0261* | 0.9901 | 0.6159 | 0.3070 |
| HippSubfield_lh_volume_GC-ML-DG-body | Regional And Tissue Volume | 0.0261* | 0.9510 | 0.2816 | 0.5469 |
| HippSubfield_lh_volume_CA4-head | Regional And Tissue Volume | 0.1672 | 0.7782 | 0.0348* | 0.9672 |
| HippSubfield_lh_volume_Whole-hippocampal-head | Regional And Tissue Volume | 0.0000*** | 0.9437 | 0.4037 | 0.4579 |
| HippSubfield_lh_volume_Whole-hippocampus | Regional And Tissue Volume | 0.0261* | 0.9548 | 0.2889 | 0.5841 |
| ThalamNuclei_lh_volume_Pc | Regional And Tissue Volume | 0.0261* | 0.9054 | 0.8823 | 0.2881 |
| aparc-Desikan_lh_thickness_fusiform | Cortical Thickness | 0.0431* | 0.9186 | 0.8564 | 0.2881 |
| aparc-Desikan_lh_thickness_parsopercularis | Cortical Thickness | 0.0753 | 0.7539 | 0.8509 | 0.0348* |
| aparc-Desikan_lh_thickness_parstriangularis | Cortical Thickness | 0.0261* | 0.9967 | 0.8540 | 0.1289 |
| aparc-Desikan_lh_thickness_precentral | Cortical Thickness | 0.0348* | 0.9777 | 0.6170 | 0.5024 |
| aparc-Desikan_lh_thickness_superiorfrontal | Cortical Thickness | 0.0431* | 0.9278 | 0.4145 | 0.7911 |
| aparc-Desikan_lh_thickness_superiortemporal | Cortical Thickness | 0.0431* | 0.8914 | 0.9081 | 0.0431* |
| aparc-Desikan_rh_thickness_parstriangularis | Cortical Thickness | 0.0000*** | 0.7998 | 0.8159 | 0.0348* |
| BA-exvivo_lh_thickness_BA6 | Cortical Thickness | 0.0348* | 0.8850 | 0.5469 | 0.7080 |
| BA-exvivo_lh_thickness_BA44 | Cortical Thickness | 0.0348* | 0.7371 | 0.7167 | 0.0000*** |
| BA-exvivo_lh_thickness_BA45 | Cortical Thickness | 0.0000*** | 0.9007 | 0.8175 | 0.1672 |
| BA-exvivo_rh_thickness_BA44 | Cortical Thickness | 0.2881 | 0.6574 | 0.9609 | 0.0261* |
| BA-exvivo_rh_thickness_BA45 | Cortical Thickness | 0.1627 | 0.5814 | 0.7676 | 0.0348* |
| aparc-DKTatlas_lh_thickness_caudalmiddlefrontal | Cortical Thickness | 0.0348* | 0.7153 | 0.5387 | 0.8551 |
| aparc-DKTatlas_lh_thickness_fusiform | Cortical Thickness | 0.0431* | 0.9800 | 0.8396 | 0.2029 |
| aparc-DKTatlas_lh_thickness_parstriangularis | Cortical Thickness | 0.0000*** | 0.9353 | 0.8321 | 0.1585 |
| aparc-DKTatlas_lh_thickness_rostralmiddlefrontal | Cortical Thickness | 0.0261* | 0.8003 | 0.9568 | 0.2971 |
| aparc-DKTatlas_lh_thickness_superiorfrontal | Cortical Thickness | 0.0431* | 0.9333 | 0.4308 | 0.7013 |
| aparc-DKTatlas_rh_thickness_parstriangularis | Cortical Thickness | 0.0348* | 0.8509 | 0.8447 | 0.0431* |
| aparc-a2009s_lh_thickness_G-front-inf-Triangul | Cortical Thickness | 0.0000*** | 0.9729 | 0.6817 | 0.0983 |
| aparc-a2009s_lh_thickness_S-front-middle | Cortical Thickness | 0.0000*** | 0.7455 | 0.8819 | 0.3293 |
| aparc-a2009s_lh_thickness_S-precentral-inf-part | Cortical Thickness | 0.0000*** | 0.9729 | 0.6289 | 0.2835 |
| aparc-a2009s_lh_thickness_S-temporal-sup | Cortical Thickness | 0.2329 | 0.9141 | 0.6734 | 0.0261* |
| aparc-a2009s_rh_thickness_G+S-cingul-Mid-Ant | Cortical Thickness | 0.4749 | 0.8212 | 0.7911 | 0.0000*** |
| aparc-a2009s_rh_thickness_G-front-inf-Triangul | Cortical Thickness | 0.0000*** | 0.9780 | 0.8175 | 0.0753 |
| aparc-a2009s_rh_thickness_S-circular-insula-inf | Cortical Thickness | 0.4661 | 0.6884 | 0.9141 | 0.0261* |
| aparc-a2009s_rh_thickness_S-circular-insula-sup | Cortical Thickness | 0.5157 | 0.8095 | 0.7911 | 0.0348* |
| aseg_global_intensity_3rd-Ventricle | Regional And Tissue Intensity | 0.0821 | 0.8867 | 0.6884 | 0.0261* |
| aseg_lh_intensity_Inf-Lat-Vent | Regional And Tissue Intensity | 0.2499 | 0.9195 | 0.7776 | 0.0000*** |

For transparency, we first report raw RMSE values (**Figure SI-6a**). As expected, across-age-group models show the largest errors, but pooled training yields slightly lower RMSE than within-age-group training. However, RMSE alone does not account for differences in target variance across test groups, making direct comparisons less meaningful. To address this, we also computed accuracy as percent improvement over a null predictor (%ΔRMSE; **Figure SI-6b**), which normalises error relative to the baseline variability of each group. Using this variance-adjusted metric, the ranking matches that observed with correlation: within-age-group > pooled > across-age-group. The gap between within and pooled models is small, whereas both clearly outperform across-age-group models. Consistent with modest effect sizes in brain-cognition prediction, absolute improvements were small (< 1%).

**Figure SI-6c** and **SI-6d** extend these analyses to the full set of training/testing combinations. Here, raw RMSE values show that the largest errors occur for *Train: Young; Test: Old*, consistent with the greater difficulty of generalising from younger to older subjects. By contrast, Train: Old; Test: Young produces relatively low RMSE despite performing poorly in correlation, likely reflecting differences in variance between test groups. To account for this, we also report %ΔRMSE (**Figure SI-6d**). This variance-adjusted measure preserves the broad ordering of within-age-group > pooled > across-age-group but highlights that *Train: Young; Test: Old* achieves only modest improvement despite good correlation. This illustrates that Pearson correlation (r) can remain high even when predictions are under-dispersed, whereas RMSE-based metrics penalise this shrinkage, leading to weaker %ΔRMSE gains.

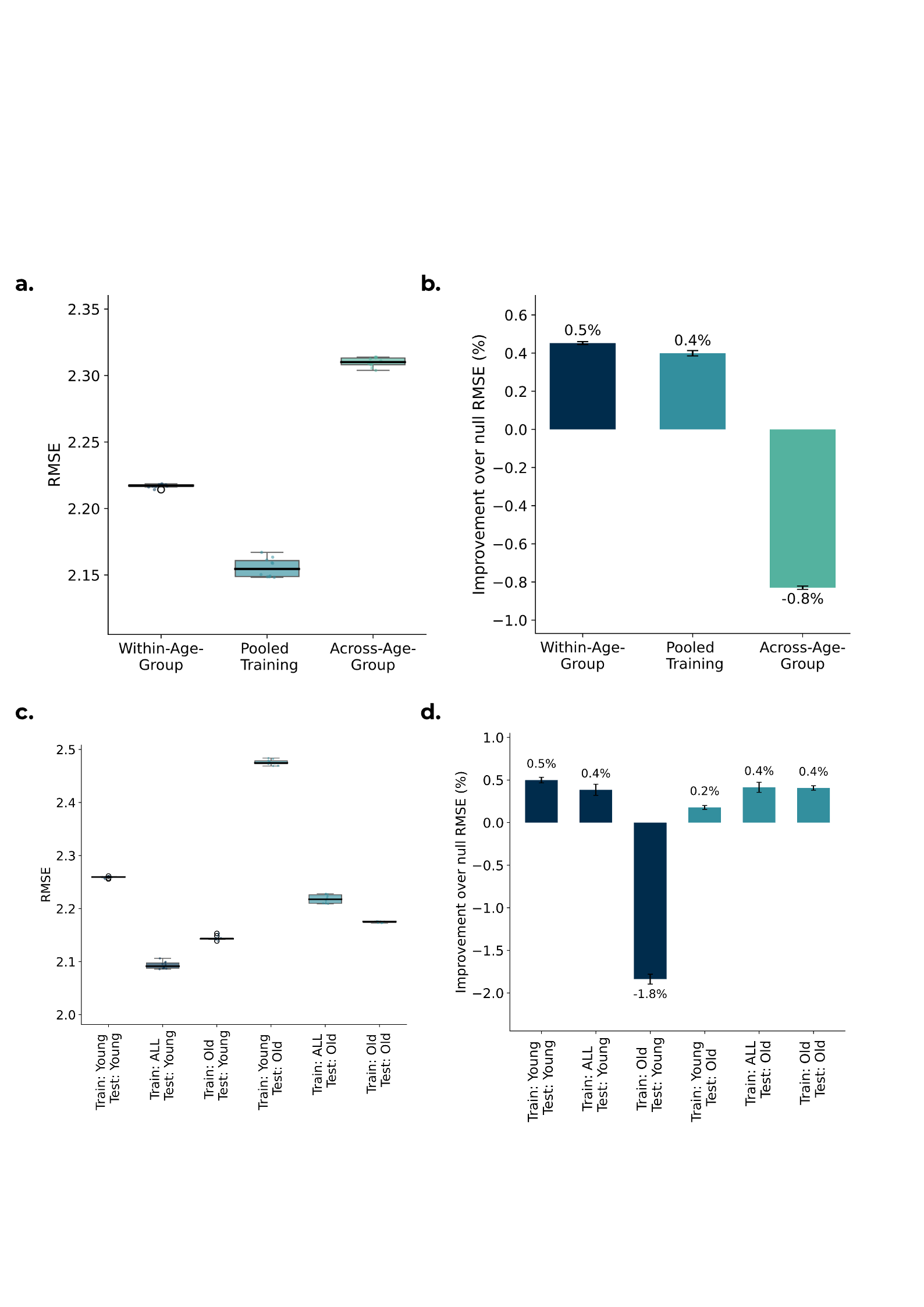

**Figure SI-6.** Model performance expressed using RMSE-based metrics. **(a)** Raw RMSE values for within-age-group, pooled, and across-age-group models. **(b)** Accuracy expressed as percent improvement over a null predictor (%ΔRMSE), where the null predictor is the test set mean of the cognitive. **(c)** Raw RMSE values for all training/testing combinations. **(d)** Corresponding %ΔRMSE values.

Elastic net is a widely used approach in neuroimaging prediction studies because it balances shrinkage and sparsity in the presence of many correlated features (Farahibozorg et al., 2021; Pervaiz et al., 2020; Roibu et al., 2023; Shen et al., 2011). We used elastic net here with both α and L1 ratio selected by nested cross-validation. In the age-specific models (Train: Young and Train: Old), the mixing parameter was consistently near zero, indicating ridge-like behaviour. This shows that differences between age-specific training strategies are driven primarily by the strength of regularisation (α), not by a shift toward sparse solutions. Pooled models (Train: ALL) selected slightly higher but still small L1 ratios (0-0.25), which may reflect the presence of features that are predictive in only one age group and are therefore pruned when training on the combined population. In line with prior work, we treat elastic net as a predictive tool rather than an inferential model. Accordingly, we do not interpret individual coefficients but instead focus on broad differences across training strategies and IDP categories.

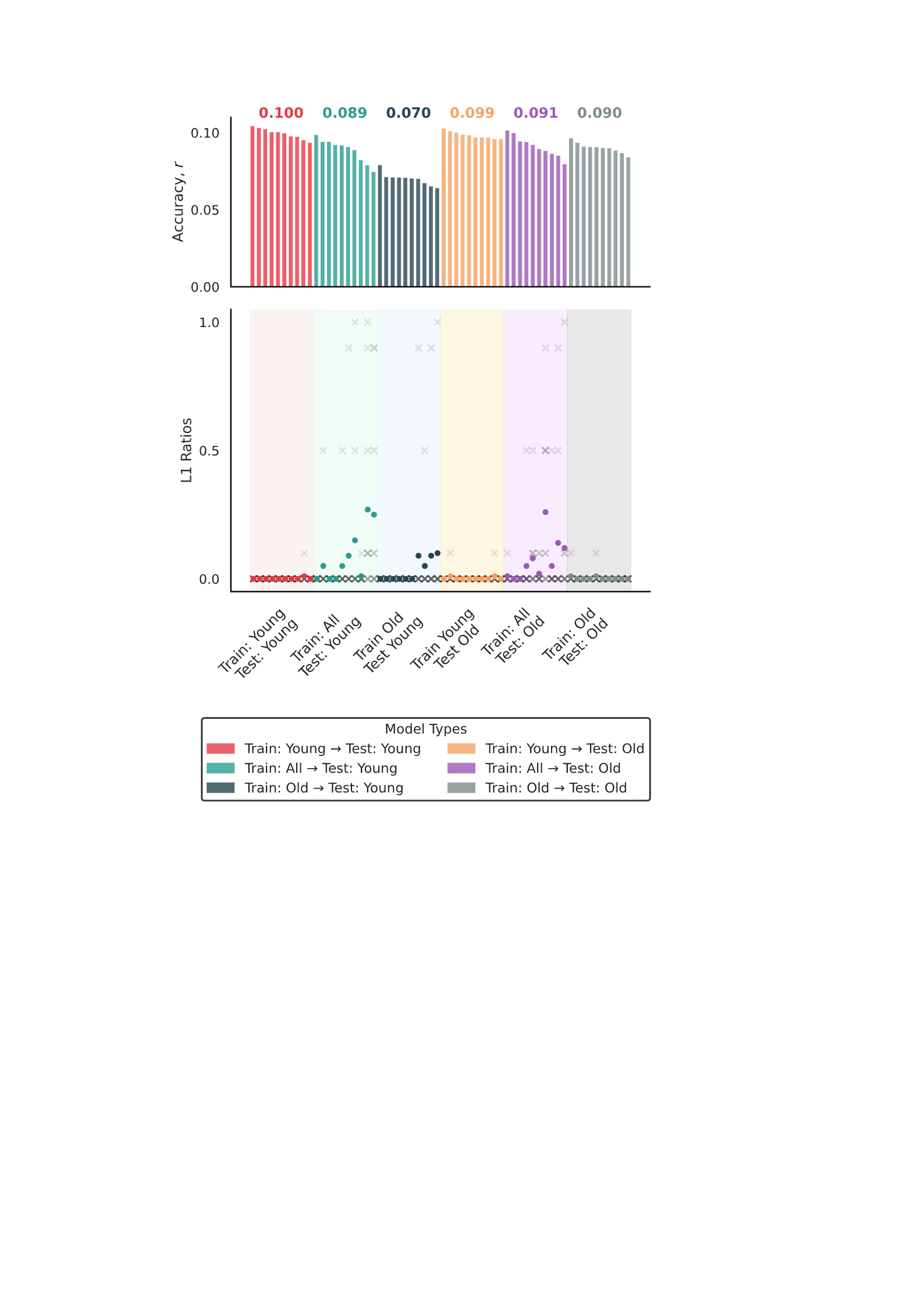

**Figure SI-7.** Elastic net mixing parameter (L1 ratio) across training strategies. **Top:** Prediction accuracy (r) sorted within each training (numbers above bars indicate the mean r). **Bottom:** L1 ratio selected by nested cross-validation for each repetition (× = individual folds; ● = mean across 10 folds). Age-specific models (Train: Young and Train: Old) clustered near L1 ratio ≈ 0, whereas pooled models (Train: ALL) consistently selected small but non-zero values. Shaded panels indicate the six training and testing conditions.
